# Pan-cancer analysis reveals complex tumor-specific alternative polyadenylation

**DOI:** 10.1101/160960

**Authors:** Zhuyi Xue, René L Warren, Ewan A Gibb, Daniel MacMillan, Johnathan Wong, Readman Chiu, S Austin Hammond, Catherine A Ennis, Abigail Hahn, Sheila Reynolds, Inanc Birol

**Affiliations:** BC Cancer Agency, Genome Sciences Centre, Vancouver, BC V5Z 4S6, Canada; Institute for Systems Biology, Seattle, 98109, USA; Department of Medical Genetics, University of British Columbia, Vancouver, BC V6H 3N1, Canada

**Keywords:** alternative polyadenylation, 3’ UTR cleavage, cancer, RNA-Seq, TCGA, *de novo* assembly, Trans-ABySS

## Abstract

Alternative polyadenylation (APA) of 3’ untranslated regions (3’ UTRs) has been implicated in cancer development. Earlier reports on APA in cancer primarily focused on 3’ UTR length modifications, and the conventional wisdom is that tumor cells preferentially express transcripts with shorter 3’ UTRs. Here, we analyzed the APA patterns of 114 genes, a select list of oncogenes and tumor suppressors, in 9,939 tumor and 729 normal tissue samples across 33 cancer types using RNA-Seq data from The Cancer Genome Atlas, and we found that the APA regulation machinery is much more complicated than what was previously thought. We report 77 cases (gene-cancer type pairs) of differential 3’ UTR cleavage patterns between normal and tumor tissues, involving 33 genes in 13 cancer types. For 15 genes, the tumor-specific cleavage patterns are recurrent across multiple cancer types. While the cleavage patterns in certain genes indicate apparent trends of 3’ UTR shortening in tumor samples, over half of the 77 cases imply 3’ UTR length change trends in cancer that are more complex than simple shortening or lengthening. This work extends the current understanding of APA regulation in cancer, and demonstrates how large volumes of RNA-seq data generated for characterizing cancer cohorts can be mined to investigate this process.

## Introduction

Alternative polyadenylation (APA) acts as a widespread and major post-transcriptional regulation mechanism for mRNAs, influencing their stabilization, translation, nuclear export, and cellular localization (Di Giammartino et al. 2011; Elkon et al. 2013; Tian and Manley 2013, 2016; Mayr 2016). APA is involved in normal cellular functions, such as cell proliferation and differentiation (Sandberg et al. 2008; Lianoglou et al. 2013; Hoffman et al. 2016), and it has also been implicated in various diseases (Ogorodnikov et al. 2016; Creemers et al. 2016; Hu et al.2016), including cancer (Singh et al. 2009; Mayr and Bartel 2009; Lin et al. 2012; Liaw et al. 2013; Xia et al. 2014; Erson-Bensan and Can 2016). Relative to normal cells, tumors have been reported to favor mRNAs with shortened 3’ untranslated regions (3’ UTRs) via APA events.Shortening may eliminate certain sequences in the 3’ UTR, such as binding sites for miRNAs and RNA-binding proteins, and other regulatory motifs, including AU- and GU-rich elements. As a result, it may enable the escape of mRNAs from degradation or other regulatory mechanisms that rely on these sequences. Therefore, APA is expected to play a role in oncogenesis, although how it contributes to the process is still poorly understood.

Standard RNA-Seq libraries have been shown to contain sufficient read evidence to identify APA events (Wang et al. 2008; Pickrell et al. 2010; Xia et al. 2014; Bonfert and Friedel 2017). Hence, collections of RNA-Seq datasets from large-scale sequencing projects like The Cancer Genome Atlas (TCGA) provide an opportunity for comprehensive investigation of APA in normal and tumor samples. Earlier, we have developed KLEAT (Birol et al. 2015), a tool that predicts cleavage sites (CSs) based on the transcripts assembled *de novo* from RNA-Seq reads, and its predictions can be used to infer APA events, because cleavage and polyadenylation of the nascent mRNA on the 3’ terminus are highly coupled reactions (Tian and Manley 2016).

Here, we characterized the CS profiles of 114 genes, mostly oncogenes and tumor suppressors (Mayr and Bartel 2009; Kumar et al. 2015), in 10,668 (9,939 tumor and 729 normal) TCGA paired-end RNA-Seq samples across 33 cancer cohorts. We identified 77 cases (i.e. gene-cancer type pairs) of differential 3’ UTR cleavage patterns between normal and tumor samples for 33 genes in 13 cancer types. For 15 of these genes, the tumor-specific cleavage patterns are recurrent in multiple cancer types. Among them, the *FGF2* gene was identified to favor a shorter 3’ UTR in tumor tissues in four cancer types (BRCA, LUAD, LUSC, and PRAD) (see Figure 1 for full names of cancer types), corroborating a previous report (Mayr and Bartel 2009). In total, we identified 16 cases of 3’ UTR shortening and 16 cases of lengthening; the remaining 45 cases indicate APA regulations that are more complex than shortening or lengthening of 3’ UTRs. Fully characterizing these cases will likely require techniques other than short read sequencing to disambiguate the relationship between individual CS predictions and multiple overlapping 3’ UTR annotations. In addition, we show that the CS profiles of this set of genes contain sufficient information to distinguish tumor samples of over 15 different cancer types, and normal from tumor samples in certain cancers.

**Figure 1.**
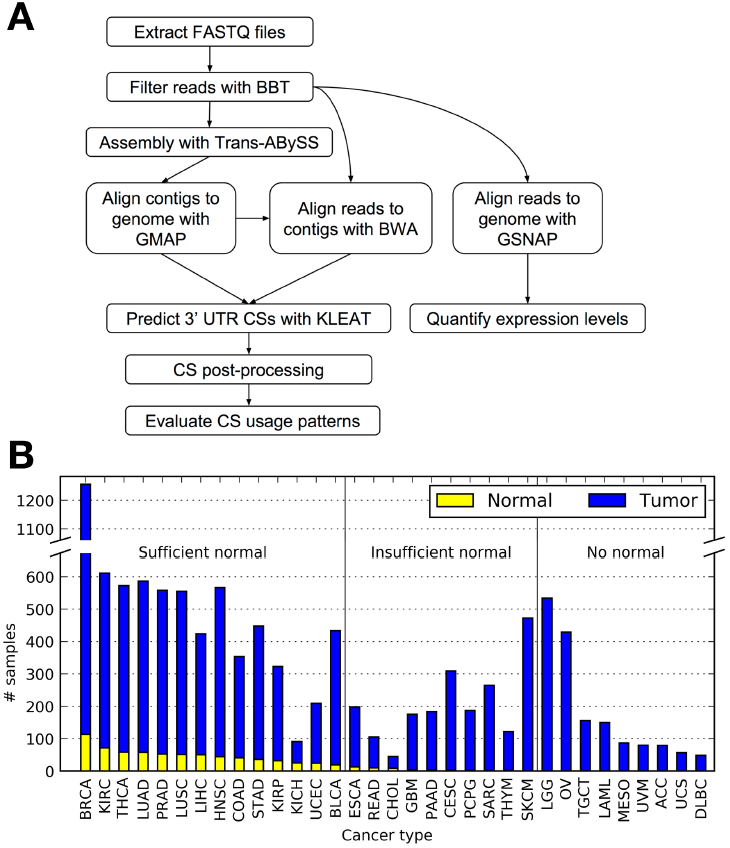
(*A*) Schematic diagram of the targeted CS prediction pipeline. (*B*) Composition of 10,668 RNA-Seq samples across 33 cancer types. TCGA RNA-Seq data are sorted in decreasing order of normal and tumor samples in each cancer. Sufficient normal means that a cancer type has ≥ 15 normal RNA-Seq samples, which is a requirement in our study to qualify for comparison between normal and tumor CS usage patterns. Insufficient normal means that a cancer type has one to 15 normal samples, and no normal means that no normal sample is available. The abbreviation of each disease is (alphabetically), ACC: adrenocortical carcinoma; BLCA: bladder urothelial carcinoma; BRCA: breast invasive carcinoma; CESC: cervical squamous cell carcinoma and endocervical adenocarcinoma; CHOL: cholangiocarcinoma; COAD: colon adenocarcinoma; DLBC: lymphoid neoplasm diffuse large B-cell lymphoma; ESCA: esophageal carcinoma; GBM: glioblastoma multiforme; HNSC: head and neck squamous cell carcinoma; KICH: kidney chromophobe; KIRC: kidney renal clear cell carcinoma; KIRP: kidney renal papillary cell carcinoma; LAML: acute myeloid leukemia; LGG: brain lower grade glioma; LIHC: liver hepatocellular carcinoma; LUAD: lung adenocarcinoma; LUSC: lung squamous cell carcinoma; MESO: mesothelioma; OV: ovarian serous cystadenocarcinoma; PAAD: pancreatic adenocarcinoma; PCPG: pheochromocytoma and paraganglioma; PRAD: prostate adenocarcinoma; READ: rectum adenocarcinoma; SARC: sarcoma; SKCM: skin cutaneous melanoma; STAD: stomach adenocarcinoma; TGCT: testicular germ cell tumors; THCA: thyroid carcinoma; THYM: thymoma; UCEC: uterine corpus endometrioid carcinoma; UCS: uterine carcinosarcoma; UVM: uveal melanoma.

## Results

### CS prediction in candidate genes

We curated a list of 114 genes, the majority of which are tumor suppressors and oncogenes (Kumar et al. 2015) (Supplemental Table S1). According to COSMIC v80 (Forbes et al. 2015), all of these genes have at least one pathogenic mutation (Supplemental Fig. S1A), and they undergo various genomic changes, such as fusion (F), mutation (M), overexpression (O), and underexpression (U), in different diseases (Supplemental Fig. S1B). Six of the genes have been reported earlier to yield transcripts with shorter 3’ UTRs in cancer cells (Mayr and Bartel 2009). To characterize the CS profiles of these genes, we developed a targeted *de novo* assembly and analysis pipeline based on KLEAT (Birol et al. 2015) (Fig. 1A). The pipeline was designed for execution on the Google Cloud Platform (GCP), a cloud computing environment. Benefiting from the scalability of the GCP, it took approximately three days to analyze 10,668 RNA-Seq samples, which consisted of 1.6 trillion reads with an average read length of 53 bp, and amounted to 67 TB of data after compression. The number of RNA-Seq samples per TCGA cancer type is shown in Figure 1B, and detailed in Supplemental Table S2.

We filtered the raw CS predictions from KLEAT to remove off-target and low-confidence predictions (Supplemental Fig. S2 and Methods) and compared the refined results to the Ensembl gene annotations (GRCh37.75) (Aken et al. 2016). We chose the last base of a 3’ UTR to represent its corresponding CS (Supplemental Fig. S3). Overall, the predicted CSs have good concordance with annotations, with 66% of them located within 15 bp of the closest annotated sites (Fig. 2A), and 79% of them having a polyadenylation signal (PAS) hexamer motif detected within a 50 bp (Xia et al. 2014) upstream window (Fig. 2B). Consistent with previous reports (Xia et al. 2014; Tian and Manley 2016; Beaudoing et al. 2000; Gruber et al.2016b), we observe a peak at around 21 bp for genes on either strand in the distribution of distances between the identified PAS hexamers and CS predictions (Fig. 2B), and we find the two most prevalent PAS hexamers are the canonical motifs AATAAA (52%) and ATTAAA (13%) (Supplemental Fig. S4). Because the precise cleavage site can vary by up to tens of base pairs (bp) (Martin et al. 2012; Tian and Graber 2012), we clustered the filtered CSs that were ≤ 20 bp apart across all samples with a single-linkage hierarchical clustering algorithm, and selected the mode coordinate within each cluster as its representative CS (Methods). We note a good concordance between representative and annotated CSs, with the distance between them spiking at 0, and note a robust distance distribution for different clustering cutoff values (Supplemental Fig. S5). Subsequent analyses are based on the representative CSs only, unless stated otherwise.

**Figure 2.**
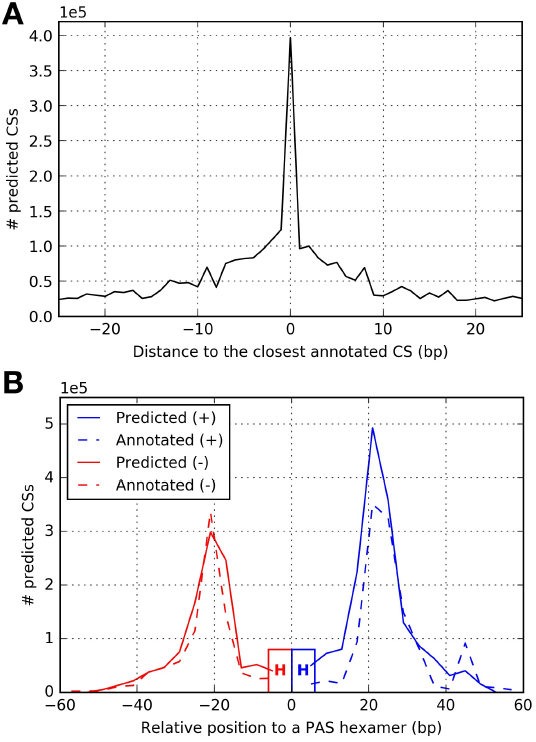
Validation of our pipeline for predicting CSs. (*A*) Distribution of the distances between a predicted CS and the closest annotated CS. (*B*) Distribution of the distances between a predicted CS and the PAS hexamer motif found within 50 bp upstream.

### Tumor-specific 3’ UTR cleavage patterns

We calculated the usage frequency of every CS in each gene in both normal and tumor samples for each cancer cohort. By comparing the CS frequencies between normal and tumor samples, we identified 77 cases of tumor-specific 3’ UTR cleavage patterns. Each case is denoted as a gene-cancer type pair, and a cleavage pattern is defined as the collection of CS frequencies within a gene in a dataset, where a dataset is defined as the normal or tumor samples from a cancer cohort. The identified 77 cases involve 33 genes across 13 cancer types. Notably, of the 14 cancer types with sufficient (≥15) normal samples available, no such pattern is found in STAD.

For 18 genes, tumor-specific cleavage patterns are observed in multiple cancer types. For 15 of these genes (*AKT2*, *BRCA1*, *CCNE1*, *CDKN2A*, *CHURC1*, *DRAM1*, *EZH2*, *FGF2*, *FGFR2*,*GNAS*, *KRAS*, *MAX*, *MITF*, *NFE2L2*, *WT1*), the tumor-specific patterns are consistent across cancer types, thus recurrent. For 6 genes (*CDKN2A*, *FLT3*, *HNF1A*, *NFE2L2*, *RNF43*, and *WT1*), the tumor-specific patterns are inconsistent across cancer types, thus disease-dependent. Three genes (*CDKN2A*, *NFE2L2*, *WT1*) are in both sets, and show recurrent patterns only in a subset of the cancer types in which they are identified.

For each case, we inferred a 3’ UTR length change trend in cancer based on the corresponding tumor-specific cleavage pattern. Here, we highlight eight cases that involve six genes (*FGF2*, *CCNE1*, *RNF43*, *CDKN2A, EZH2*, and *PTCH1)*. The first four display 3’ UTR shortening or lengthening trends in tumors (Fig. 3), and the other four show more complex cleavage patterns that do not fit the shortening/lengthening paradigm (Fig. 4). All other cases are depicted in Supplemental Figure S6. Gene-level and CS-level summaries for all 77 cases are provided in Supplemental Table S3 and S4, respectively.

**Figure 3.**
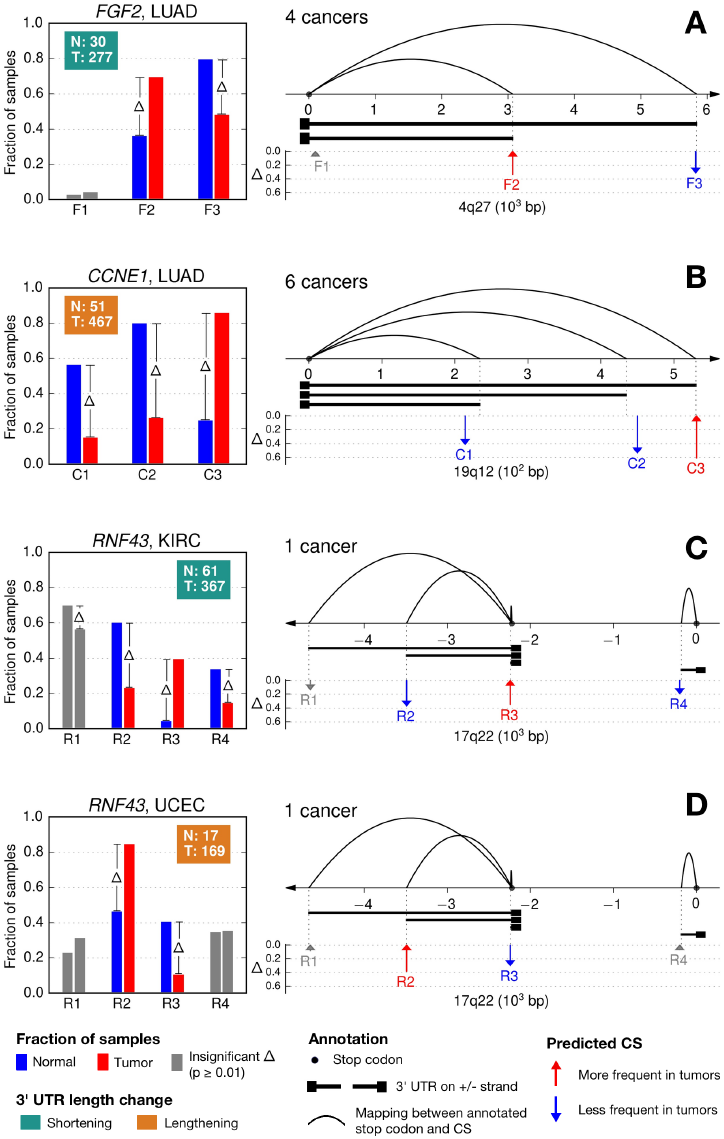
Three representative genes showing tumor-specific CS usage patterns in different cancer types, with implications on their 3’ UTR length changes. (*A*) *FGF2*. (*B*) *CCNE1*. (*C*,*D*) *RNF43*. (*A*-*D*) Inside each panel, to the left, the bar plot shows the usage frequencies of CSs within a given gene. Each group of bars represents the frequencies of one CS in normal (blue) and tumor (red) samples, respectively. Bar groups are ordered by the genomic coordinates of their corresponding CSs, and those corresponding to insignificant (p ≥ 0.01, Fisher’s exact test) frequency change are greyed out. Bar width is not meaningful. The text box shows the number of normal (N) and tumor (T) samples considered for frequency calculation with sufficient gene expression. The background of the text box encodes the trend of 3’ UTR length change with respect to the CS usage patterns in normal. To the right, the number of cancers supporting recurrent tumor-specific CS usage patterns are shown beside the bar plots. The arcs represent mappings between annotated stop codons and CSs with corresponding 3’UTRs drawn below. 3’ UTRs drawn here may also contain introns. Vertical arrows indicate the positions of predicted CSs by KLEAT. Annotated and predicted CSs match well, but are not expected to overlap exactly. An arrow pointing upwards or downwards represents an increase (red) or decrease (blue) in frequency from normal to tumor, respectively, with its height representing the difference (Δ). Like bars, arrows are greyed out if the corresponding difference is insignificant. The X-axis represents the genomic coordinates offset by the position of the first stop codon, and a horizontal arrow indicates the gene strand. For clarity, CSs with usage frequencies lower than 5% in both normal and tumor samples, and that do not undergo any significant change in any cancer type considered herein are not shown. For a comprehensive view of all CSs with distribution of gene expression levels, see Supplemental Figure S6.

**Figure 4.**
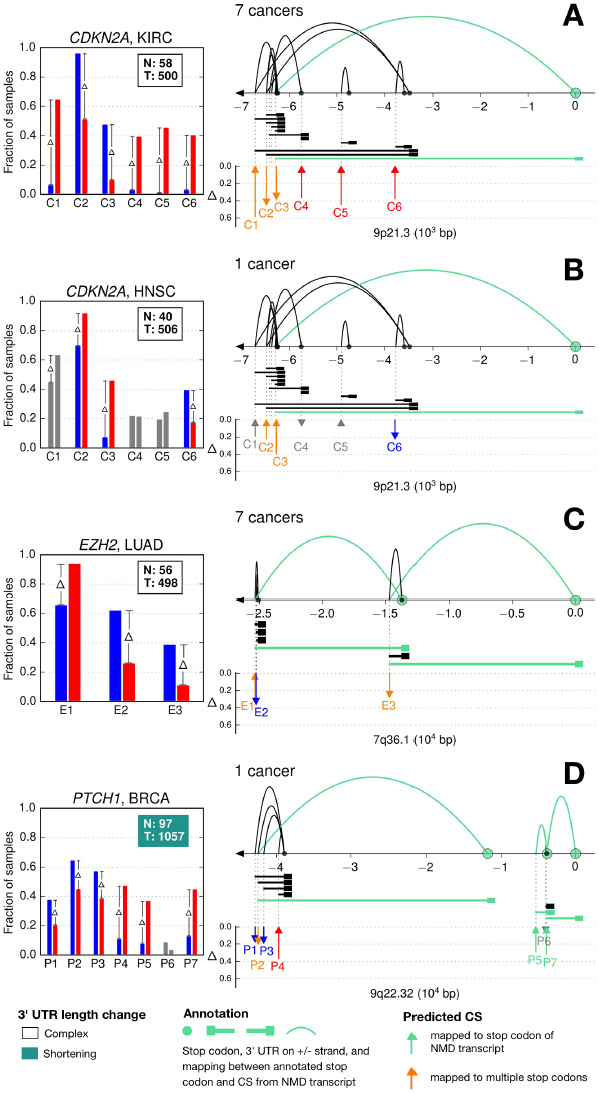
Three representative genes showing complex tumor-specific CS usage patterns in different cancer types. (*A*,*B*) *CDKN2A*. (*C*) *EZH2*. (*D*) *PTCH1*. (*A*-*D*) The legend of Figure 3 applies. In addition, when the 3’ UTR length change is too complex to be resolved into a shortening or lengthening trend, the corresponding text box is left uncolored. NMD-related transcript elements are colored in cyan, as are the corresponding arrows for predicted CSs. An orange arrow indicates that a predicted CS that undergoes significant usage frequency change is mapped to multiple stop codons, with its associated 3’ UTR length being ambiguous.

The *FGF2* gene (Fig. 3A) presents an example of 3’ UTR shortening that has been previously reported in several cancer cell lines (Mayr and Bartel 2009). The predicted and annotated CSs have strong concordance, but their coordinates are not expected to match exactly. We use a convention that labels CSs by a letter from the gene name followed by an index incremented relative to the CS genomic coordinates on the positive strand, so the three predicted CSs of *FGF2* are marked as F1, F2, and F3. For *FGF2*, a positive strand gene with a single annotated stop codon, an increment in the index (e.g. F2 to F3) indicates an increase in the 3’ UTR length. The F1 site is over 2 kb from the closest annotated CS, and demonstrates the ability of our analysis to detect potential novel CSs. F1 has a low frequency of usage in both normal and tumor samples (typically less than 20%, Supplemental Table S4), and its frequency does not undergo any significant change (p > 0.01, Fisher’s exact test) from normal to tumor. In contrast, F2 and F3 are frequently used. In four TCGA cohorts (LUAD, BRCA, LUSC, and PRAD), the frequency of F2 increases in tumor samples, while that of F3 decreases, both significantly (p = 0.0004 and 0.001, respectively). Therefore, it indicates that FGF2 undergoes 3’ UTR shortening in the tumor tissues that are associated with these cancer types.

*CCNE1* (Fig. 3B) presents an example of 3’ UTR lengthening in the tumor tissues of six cancer cohorts (LUAD, BRCA, HNSC, KIRP, LIHC, and LUSC). It also has a single annotated stop codon, and its two shorter 3’ UTR isoforms, associated with CSs labeled C1 and C2, decrease in frequency in tumor samples, while a longer form associated with C3 has an increased frequency, thus indicates 3’UTR lengthening.

In contrast to *FGF2* and *CCNE1*, the *RNF43* gene (Fig. 3C,D) has two annotated stop codons, and it shows both 3’ UTR shortening and lengthening trends, depending on the tissue of origin. *RNF43* has four predicted CSs. Since it is a negative strand gene, for the three CSs (R1, R2, and R3) that share the same stop codon, an increment in the index (e.g. R2 to R3) indicates a decrease in the 3’ UTR length. Among them, R2 and R3 experience significant shifts in their frequencies from normal to tumor in both KIRC and UCEC. Intriguingly, these shifts occur in opposite directions in the two diseases, indicating 3’ UTR shortening in KIRC and lengthening in UCEC. As for R4, which is associated with a different stop codon, we also note a significant usage frequency shift in KIRC, indicating biological condition specific expression of corresponding alternative transcript isoforms.

Unlike these clear and simple cases of shortening or lengthening, most cases we observed show more complex APA regulations. The *CDKN2A* gene, like *RNF43*, also displays multiple disease-dependent frequency shifts. Of its six predicted CSs, three (C2, C3, and C6) display significant frequency changes in KIRC and HNSC (Fig. 4A,B). Both C2 and C3 decrease in frequency in the former cancer but increase in the latter, while C6 shows the opposite change. However, *CDKN2A* has seven annotated stop codons, many more than the genes described above, and some of its CSs exhibit one-to-many relationships to its stop codons. Specifically, C1, C2, and C3 (highlighted in orange) can be associated with multiple stop codons; thus, the frequencies of their associated 3’UTR lengths cannot be determined unambiguously. On the other hand, C4, C5, and C6 match 3’ UTRs that have no overlap; thus, comparing the length of their matched UTRs is not straightforward. Non-overlapping 3’ UTRs indicate transcript isoforms with different translated gene products, hence their frequency shifts in KIRC potentially relate to the preferential usage of these isoforms in normal and tumor kidney tissues. In addition to the above complexity, one of the stop codons (on the annotation matched to C3) is involved in nonsense mediated decay (NMD), which is a surveillance mechanism for removing prematurely transcribed mRNAs (Lykke-Andersen and Jensen 2015; Lindeboom et al. 2016). We observe that NMD transcripts tend to have longer 3’ UTRs compared to the protein coding transcripts (P=6.5×10^−16^,Kolmogorov–Smirnov test) (Supplemental Fig. S7), but the meaning of such a comparison is not clear, as the former may not result in any translated polypeptide. The intricate tumor-specific 3’ UTR cleavage pattern of C*DKN2A* in KIRC is also consistently identified in six other cancer types including COAD, KICH, KIRP, LIHC, PRAD and THCA.

Much like *CDKN2A* in KIRC, the *EZH2* gene (Fig. 4C) also displays a recurring and consistent tumor-specific cleavage pattern in seven cancer types (BRCA, KIRC, KIRP, LIHC, LUAD, PRAD, THCA). In addition, it illustrates another level of complexity, as its second stop codon (near E3) is shared by both protein coding and NMD transcripts. Thus, *EZH2* presents a many-to-many-to-many relationship among CSs, stop codons, and transcript types.

Finally, we show the *PTCH1* gene (Fig. 4D) as an example of 3’ UTR length change trend that is resolvable despite complex CS frequency shifts. During the resolution process (Methods), we ignored P5, P6, and P7 because they were each mapped to a separate stop codon, and we also ignored P2 since it was mapped to multiple stop codons and hence associated with an ambiguous 3’ UTR length. For the remaining CSs, P1 and P3 correspond to longer 3’ UTRs, and their usage frequencies decrease in tumors, while P4 is associated with a shorter 3’ UTR, and its usage frequency increases in tumors. Therefore, in BRCA, *PTCH1* could be tentatively considered to display a 3’ UTR shortening trend.

In total, we investigated the 3’ UTR length change trends of all 77 cases of tumor-specific cleavage patterns by conducting stop codon-level comparisons (Fig. 5 and Methods). We identified a comparable number of 3’ UTR shortening (n=16) and lengthening cases (n=16), with the remaining 45 cases labeled as complex trends.

**Figure 5.**
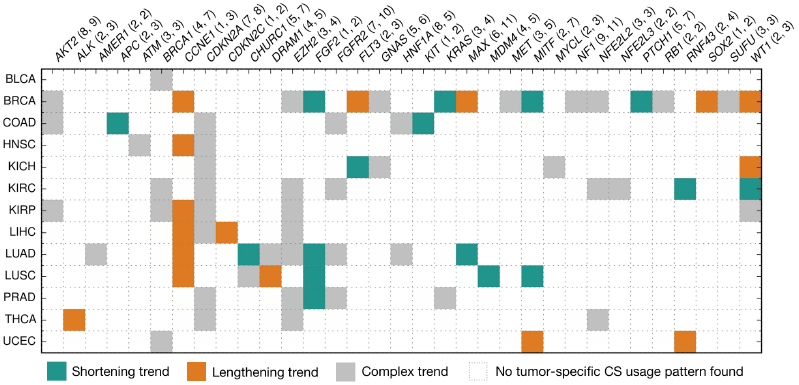
3’ UTR length modulation trends for all 77 reported cases with tumor-specific CS usage patterns identified. The numbers in parentheses indicate the numbers of annotated stop codons and CSs per gene. For example, *AKT2* (8, 9) means the gene has 8 annotated stop codons and 9 annotated CSs.

### CS profiles distinguish tissues and biological states

Presence of CSs is conditional on the expression of the transcripts they originate from, which may be tissue and/or biological condition specific. We also explored the utility of CS profiles in distinguishing cancer cohorts and the disease states of samples. To do that, we encoded CS profiles of samples as binary vectors marking presence or absence of CSs, and visualized those vectors in a 2-dimensional space using t-Stochastic Neighbor Embedding (t-SNE) (Platzer 2013).

t-SNE is a nonlinear dimensionality reduction method that retains local structures in a dataset, ignoring global structures. This means, in a reduced space samples with similar CS profiles are expected to form clusters that are separated by nondeterministic inter-cluster distances.

Using t-SNE, we first visualized the CS profiles of 9,939 tumor samples from all 33 cancer types. Exploring our dataset in 2-dimensional embeddings, we observed that 17 out of 33 cancer types were more co-localized (Supplemental Fig. S8), so we reapplied t-SNE on these 17 cancer types (Fig. 6A). Overall, CS profiles distinguish tumor samples of different cancer types, including two from the same organ – LGG and GBM, two brain cancers. In contrast, two cancer types from the digestive tract, ESCA and STAD, form two overlapping clusters, suggesting that they have similar CS profiles.

**Figure 6.**
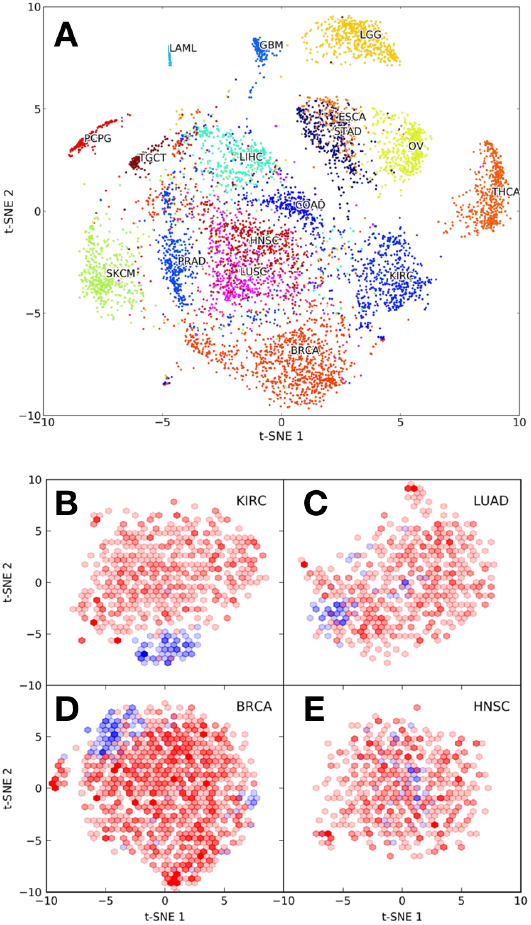
CS profiles distinguish cancer types and sample types. (*A*) t-SNE visualization of the CS profiles of 7,105 tumor samples from 17 cancer types. (*B*-*E*) t-SNE visualization of the CS profiles of normal (blue) and tumor samples (red) per cancer type for KIRC, LUAD, BRCA and HNSC. t-SNE coordinates are transformed into 2-dimensional histograms and shown in hexagon bins, with color intensity indicating the count in a bin.

We then applied t-SNE to normal and tumor samples from 14 cancer cohorts with at least 15 normal samples. Figure 6B-E shows four cancer types with different levels of separation between their normal (blue) and tumor (red) samples, with the rest plotted in Supplemental Figure S9. For certain cancer types such as KIRC, LUAD, and BRCA (Fig. 6B-D), the normal samples co-localize, implying that their CS profiles have characteristic differences from those of tumor samples. In HNSC, however, such co-localization is less pronounced, suggesting that CS profiles are similar in these tumor and normal samples (Fig. 6E).

## Discussion

We have profiled the CSs of 114 genes in 10,668 TCGA RNA-Seq samples using a targeted analysis pipeline based on KLEAT (Birol et al. 2015). By comparing the frequencies of predicted CSs between normal and tumor samples, we identified 77 cases of tumor-specific 3’ UTR cleavage patterns involving 33 genes across 13 cancer types. For 15 genes, the tumor-specific patterns are recurrent in multiple cancer types, consistent with a previous report on recurrence of tumor-specific APA events (Xia et al. 2014). However, for certain genes (e.g. *RNF43*,*CDKN2A*), their tumor-specific cleavage patterns, even when observed in multiple diseases, may manifest in a disease-dependent manner. Specifically, while the frequency of a CS increases in tumors in one cancer type, it may decrease in another. Such patterns imply recurrent and disease-specific oncogenic mechanisms in cancers of different tissues of origin. In contrast to a two-polyA-site model (Xia et al. 2014), 40% (31/77) of the cases reported here have more than two CSs that undergo significant frequency changes (p < 0.01, Fisher’s exact test).

In general, we expect many more cases of differential 3’ UTR cleavage patterns between tumor and normal samples than what is reported here. Note that, in our study we enforced the co-occurrence of both a significant increase and a significant decrease in sample frequency for at least two separate CSs within the same gene (Methods). We also limited the scope of our study to 114 genes.

Although previous reports have emphasized the shortening of 3’ UTRs in cancer cells (Mayr and Bartel 2009; Xia et al. 2014), our analysis suggests that lengthening may be equally frequent. Lengthening of 3’ UTRs has also been reported in other APA studies, such as those on breast(Fu et al. 2011) and colorectal (Morris et al. 2012) cancers, but its roles in cancer biology remain unclear. In principle, while shortening may enable escape from inhibition of gene expression, conceivably promoting unchecked growth, lengthening may potentiate repression of anti-proliferative genes_both having the potential to abet cancer progression. For certain genes(e.g. *RNF43*), 3’ UTR length changes could be disease-dependent, with shortening in one cancer, but lengthening in another.

For many genes, even when differential 3’ UTR cleavage patterns between normal and tumor tissues are identified, it may be difficult to resolve them into a simplistic shortening or lengthening pattern at the isoform level, especially when a CS is associated with multiple stop codons. In the results reported here, 33/297 (11%) of the predicted CSs in 33 genes are mapped to more than one stop codon. In such cases, it is not possible to quantify the usage of corresponding 3’ UTRs based on CS predictions alone, other than providing an aggregate measure of all the 3’ UTRs cleaved at the same genomic coordinates. In general, genes with multiple stop codons are very common, and account for 28 out of 33 genes with tumor-specific cleavage patterns reported here, or about 68% of all human protein coding genes based on the Ensembl annotations used. Even when the CS-to-stop codon mappings are unambiguous, if multiple 3’ UTRs are associated with different stop codons, and have no or limited overlap, their interpretation within the old paradigm of length modifications of 3’ UTRs may be difficult, as is the case for the *CDKN2A* gene. In addition, about half (16/33) of the reported genes have stop codons related to NMD transcripts (Lykke-Andersen and Jensen 2015; Lindeboom et al. 2016), and in some genes (e.g. *EZH2*), a stop codon can be used in both protein coding and NMD transcripts.

Taking advantage of TCGA RNA-Seq data, we demonstrated that retrospective analysis can add value to publicly available data from large cohort studies. Our results extend the current understanding of APA regulation in cancer, revealing a more complex pattern than simple 3’UTR shortening or lengthening. We also demonstrated that CS profiles of samples might have enough information to cluster samples with respect to their tissues of origin, and even differentiate certain cancer types from common tissue types.

## Methods

### Collection of RNA-Seq data and metadata

We used a copy of the TCGA RNA-Seq data hosted by the Institute for Systems Biology-Cancer Genomics Cloud (ISB-CGC) pilot on the Google Cloud Storage, part of the Google Cloud Platform (GCP). Most of the metadata were extracted from a MANIFEST file from the CGHub (Wilks et al. 2014). In total, 10,668 samples were analyzed, with each sample identified by a unique analysis ID. All analysis IDs are listed in Supplemental Table S5. Sample types were generalized as ‘normal’ (solid tissue normal) and ‘tumor’ (which includes primary solid tumor, metastatic, recurrent solid tumor, additional - new primary). A more detailed description of the protocols for data collection is provided in the Supplemental Methods.

### Curation of genes for analysis

Of the genes studied, 99 have been reported to be either an oncogene or a tumor suppressor (Kumar et al. 2015), six have been reported to favor transcripts with shorter 3’ UTRs in cancer (Mayr and Bartel 2009), and 10 were from in-house studies. *DICER1* is in both of the first two sets, bringing the total to 114 genes.

### Design of the targeted CS prediction pipeline

RNA-Seq reads were first filtered against the candidate genes using the biobloomcategorizer utility from BioBloomTools (BBT) (Chu et al. 2014). The resulting categorized reads were then assembled into contigs with Trans-ABySS (Robertson et al. 2010), and these contigs were in turn aligned to the reference human genome with GMAP (Wu and Watanabe 2005). The raw reads were aligned to both the assembled contigs with BWA (Li and Durbin 2009), and the reference genome with GSNAP (Wu and Nacu 2010). Both contig-to-genome and read-to-contig alignment results were used for CS prediction by KLEAT (Birol et al. 2015), and the read-to-genome alignments were used for both expression level quantification and assessment of KLEAT predictions.

### Implementation of the pipeline

The pipeline was implemented in Python with the Ruffus framework (Goodstadt 2010). The software used include SAMtools-0.1.18 (Li et al. 2009), BioBloom tools-2.0.12 (Chu et al. 2014), Trans-ABySS-1.5.2 (Robertson et al. 2010) and ABySS-1.5.2 (Simpson et al. 2009), GMAP- 2014-12-28 (Wu and Watanabe 2005; Wu and Nacu 2010), and BWA-0.7.12 (Li and Durbin 2009). The source code also includes a copy of the specific version of KLEAT.py used in this study. For its use on the GCP, a Docker image of the pipeline can be built from the Dockerfile included in the source code.

### Execution of the pipeline

We executed the pipeline on the ISB-CGC powered by the GCP. For each RNA-Seq sample, a virtual machine (VM) instance with four vCPUs, 20 GB of memory, and a sample size-dependent amount of persistent disk was used. For each instance, we requested a sufficient disk size for storing both input data and intermediate and final results, calculated as 30 × Size(sample) + 50 GB. The scaling factor of 30 is based on experience in pilot runs, and the extra 50 GB was reserved for storing reference data. Google Genomics Pipelines API (https://cloud.google.com/genomics/reference/rest/v1alpha2/pipelines) was used to orchestrate all VM instance tasks including creation, deletion, and data transfer, and it substantially reduced the administrative workload.

The reference data included an hg19 reference genome (Lander et al. 2001), the GMAP/GSNAP (Wu and Watanabe 2005; Wu and Nacu 2010) index of hg19, a prebuilt BioBloom filter (Chu et al. 2014) of all the candidate genes’ transcripts, and a specific version of the gene annotation used by KLEAT (Birol et al. 2015) (more details on annotations are available in Supplemental Methods).

The BioBloom filter was built with the biobloommaker utility from BBT (Chu et al. 2014). As for the input to biobloommaker, all transcripts of all the candidate genes from the Ensembl annotation (Aken et al. 2016) were used. The annotated sequences were augmented by 300 bp flanking sequences on both ends of each transcript to collect RNA-Seq reads that were partially aligned to them.

During the *de novo* assembly of transcripts for each sample, three *k*-mer sizes were used, depending on the corresponding read length: (22, 32, 42), (32, 52, 72), and (32, 62, 92) were used for samples with read lengths of 45-50, 75-76, and 100 bp, respectively.

### Annotation pre-processing

The Ensembl annotation was downloaded from http://ftp.ensembl.org/pub/release-75/gtf/homo_sapiens/Homo_sapiens.GRCh37.75.gtf.gz, and then pre-processed before being used. First, we extracted the annotated CSs of all protein coding and NMD transcripts that were CDS 3’ complete (without cds_end_NF tag, https://www.gencodegenes.org/gencode_tags.html) for all candidate genes. To calculate 3’ UTR lengths, we also extracted the mapping information between annotated CSs and stop codons from transcripts. A more detailed description of the extraction process can be found in Supplemental Methods. After extraction, we discarded transcript-level information, since a predicted CS is transcript agnostic, and removed redundant mapping relationships caused by multiple transcripts sharing the same CS and stop codon.Lastly, we clustered the annotated CSs as described in **CS Clustering**.

### CS prediction and post-processing

The CSs were predicted by KLEAT with all parameters set to default. We post-processed the KLEAT results before any CS usage analysis (Supplemental Fig. S2). Specifically, we used the following columns output by KLEAT: gene, transcript_strand, chromosome, cleavage_site, length_of_tail_in_contig, number_of_bridge_reads, and max_bridge_read_tail_length.

In total, 67,544,140 CSs were predicted across 10,668 samples. First, we filtered out off-target CSs by only selecting those that were associated with the candidate genes, keeping 17% of the predictions. After initial filtering by genes, we reassigned each remaining CS to the closest clustered annotated CS (See **Annotation pre-processing**), and then calculated the signed distance between them. We also calculated the location of PAS hexamer motifs, if present, searching up to a 50 bp window upstream of a predicted CS. When multiple PAS hexamers existed in the window, the strongest one was picked (Beaudoing et al. 2000). Next, we applied another filter to select the most confident predictions. Specifically, a predicted CS must meet at least one of the following two criteria to be retained: 1) Its distance to the closest annotated CS was required to be 25 bp or less. The 25-bp threshold was chosen by considering the distance distribution shown in Supplemental Fig. S10, and taking a threshold at the plateau. This criterion was designed for selecting CSs that had already been annotated. 2) One of the two strongest PAS hexamers AATAAA and ATTAAA (Gruber et al. 2016a) were required to be within a 50 bp window, and at least one of the following conditions of polyadenylation evidence was satisfied: length_of_tail_in_contig ≥ 4, number_of_bridge_reads ≥ 2, or max_bridge_read_tail_length ≥ 4. The second criterion is an empirical one that is independent of annotation, and it is designed mainly for selecting potential novel CSs. We verified that AATAAA and ATTAAA were the two most frequent PAS hexamers associated with the predicted CSs both before and after the second filtering steps (Supplemental Fig. S4). After the two filtering steps, about 5% of the CSs were retained and clustered as described in **CS Clustering**. The post-processing steps resulted in 2,136 unique predicted CSs in 114 target genes across all samples.

### CS clustering

We used the single-linkage hierarchical clustering algorithm to combine CSs that were ≤ 20 bp apart, iterating when necessary for clusters to converge. After clustering, we selected the mode CS coordinate within each cluster as its representative location. If multiple modes existed, the median of the modes was used. Then, every CS was associated with one of the representative CSs, and multiple CSs associated with the same representative CS were merged within each sample. The clustering method was independently applied to both annotated and predicted CSs.

The clustering process inevitably decreases the detection resolution, so our analysis is not able to distinguish CSs that were closer than the clustering cutoff (20 bp). However, the clustering results were insensitive to different cutoff values, albeit the number of clusters would vary (Supplemental Fig. S5).

### CS usage frequency calculation

For a given CS in a gene, its usage frequency was calculated as the fraction of normal or tumor samples that were predicted to use it. For each sample, we required the gene under consideration to have at least one CS predicted; otherwise, its expression was considered insufficient, and the sample was excluded from the frequency calculation.

### Comparison of cleavage patterns between normal and tumor samples

For every predicted CS of every gene in each cancer type, we calculated usage frequencies in normal and tumor samples, and then evaluated the significance of the frequency difference with a Fisher’s exact test. The input to the test included the number of normal and tumor samples with and without a CS. The frequencies of multiple CSs within one gene collectively formed a cleavage pattern for that gene, and to report difference in patterns between normal and tumor, we required the co-occurrence of at least one significant increase (p < 0.01) in the frequency of one CS, and at least one significant decrease in that of another CS. Without this requirement, an increase or decrease in the CS frequency could simply be a result of gene up- or down-regulation in the normal or tumor samples. For example, if a gene is down-regulated in tumor, the decreases in the usage frequencies of certain CSs was expected due to a lower probability of CS prediction by KLEAT (Supplemental Fig. S11). However, when the increase in the use of other CSs was required, it implied involvement of regulation mechanisms of APA events (Di Giammartino et al. 2011; Elkon et al. 2013; Tian and Manley 2013, 2016).

In addition, to justify the comparison of CS usage frequencies between two groups of samples, we also verified that the RNA-seq data for normal and tumor samples in TCGA have comparable sequencing depths (Supplemental Fig. S12). For CS predictions, comparable sequencing depths serve to normalize data among samples, as different sequencing depths yield different CS prediction probabilities.

### Resolution of the 3’ UTR length change trends

We first mapped a predicted CS to the closest annotated one. If it was >25 bp away, the predicted CS was considered potentially novel, and was ignored for length trend resolution. The 25-bp cutoff was selected after trying a range of values (Supplemental Fig. S13). After mapping, we determined the associated stop codons for each CS. If a CS was associated with only a single stop codon, its corresponding 3’ UTR length was unambiguously calculated and used for trend resolution. All CSs mapped to multiple stop codons were ignored. A detailed description of the trend resolution approach is provided in Supplemental Methods.

### t-SNE transformation

A CS profile is encoded in a binary vector using the set of all representative CSs, with 1 indicating a prediction and 0 meaning no prediction. First, high-dimensional CS profile data were reduced to 50 dimensions with truncated singular value decomposition (SVD), and then further reduced to 2 dimensions with t-SNE. Different settings of the t-SNE parameters perplexity and learning rate were used, depending on the sample size, as follows. When transforming all tumor samples from 33 or 17 cancer types, perplexity (PX) was set to 50, and learning rate (LR) was set to 1000. In general, for the sample sizes used, the performance of t-SNE was observed to be qualitatively robust to these parameters (Van Der Maaten and Hinton 2008). However, when transforming normal and tumor samples for a single cancer type, PX and LR were lowered to account for a much smaller sample size in the cohorts. For BRCA, which has the largest number of samples of all the cancer types, we used PX=40 and LR=500. For cohorts with the smallest number of samples above our cutoff of 15, KICH and UCEC, we used PX=15 and 25, and LR=30 and 100, respectively. For other cancer types, we used PX=40 and LR=300. When visualizing normal and tumor samples of the same cancer type, we removed the means from the two t-SNE dimensions to center them at the origin, and then transformed the coordinates into a 2-dimensional histogram with hexagon bins.

### Python libraries used

In addition to the aforementioned Ruffus framework (Goodstadt 2010), we used several other Python libraries for scientific computing (Oliphant 2007) to facilitate our analysis. The t-SNE implementation of Scikit-learn-0.18 (Pedregosa et al. 2011) was used to transform CS profiles. The hierarchical clustering algorithm implemented in SciPy-0.18.1 (Jones et al. 2001; van der Walt et al. 2011) was used for CS clustering. Pandas-0.19.0 (Mckinney 2009) was used for tabular data transformation and analysis. Matplotlib-1.5.3 (Hunter 2007) was used for plotting.Jupyter-1.0.0 notebook (Perez and Granger 2007) was used for tracking analysis steps and results.

### Data access

The curation process for candidate genes and their transcript sequences for BioBloom filter generation is documented in detail at https://github.com/bcgsc/utrtargets. The source code of the implemented targeted CS prediction pipeline is available at https://github.com/bcgsc/tasrkleat. A copy of all the reference and metadata used is provided at http://bcgsc.ca/downloads/tasrkleat-static/.

## Acknowledgements

We thank Dr. A. Gordon Robertson for his suggestions on figures and writing, and Dr. Erin Pleasance for discussions on oncogenes and tumor suppressor genes. This work has been supported in part by the National Human Genome Research Institute of the National Institutes of Health under award number R21CA187910. Additional support was provided by the Canadian Institutes of Health Research. The high throughput analysis in this study was performed on the Institute for Systems Biology-Cancer Genomics Cloud (ISB-CGC), a pilot project of the National Cancer Institute (under contract number HHSN261201400007C). The content of this manuscript is solely the responsibility of the authors, and does not necessarily represent the official views of the National Institutes of Health or other funding organizations. The results shown here are in part based upon data generated by the TCGA Research Network: http://cancergenome.nih.gov/.

## Author contributions

Z.X. designed and performed the data collection and analysis. Z.X., R.L.W., and I.B. interpreted the results and wrote the manuscript. R.L.W. and Z.X. designed the targeted CS prediction pipeline. Z.X. implemented the pipeline. E.A.G. and R.C. collected the candidate genes. J.W. and Z.X. curated mutational information of the genes. R.C. and D.M.developed KLEAT. Z.X., J.W. and R.C. optimized the CS filters. S.A.H. contributed to annotation, data pre-processing and writing the manuscript. S.R. and A.H. conducted FastQC analysis for all RNA-Seq samples on the GCP. C.A.E. provided project management support.R.L.W. and I.B. conceived and supervised the study.

